# Hepatic Ketogenesis Attenuates Cardiac Hypertrophy via Metabolic Reprogramming

**DOI:** 10.64898/2026.01.30.702687

**Authors:** Toshihiro Yamada, Kei Morikawa, Akira Fujiyama, Takumi Nagakura, Yuqing Xu, Miho Kataoka, Terumasa Umemoto, Miki Bundo, Kazuya Iwamoto, Zeeshan Ahmed, Suvi Linna-Kuosmanen, Shinsuke Hanatani, Hiroki Usuku, Yasushi Matsuzawa, Yasuhiro Izumiya, Eiichiro Yamamoto, Kenichi Tsujita, Yuichiro Arima

## Abstract

**Background:** Heart failure with preserved ejection fraction (HfpEF) is increasingly recognized as a multisystem disorder linked to the cardiovascular-kidney-metabolic (CKM) syndrome. While the falling heart undergoes metabolic reprogramming, the interorgan crosstalk regulating myocardial substrate preference in HFpEF remains elusive. We aimed to clarify the role of systemic and local ketogenesis in the pathogenesis of cardiac hypertrophy and HFpEF.

**Methods:** A mouse model of HFpEF was employed using a high-fat diet combined with NG-Nitro-L-arginine methyl ester hydrochloride (L-NAME). Cardiac hypertrophy and systemic metabolic profiling including ketogenesis were evaluated. To dissect the role of site-specific ketogenesis, we generated inducible cardiomyocyte-specific (*Hmgcs2^ΔiCM^*) and hepatocyte-specific (*Hmgcs2^ΔHep^*) knockout mice of HMG-CoA synthase 2 (*Hmgcs2*), deficient in the rate-limiting enzyme for ketogenesis. Cardiomyocyte -specific nuclei were isolated for transcriptomic (RNA-seq) and *in vitro* assays in H9C2 cells were used to elucidate molecular mechanisms.

**Results:** The HFpEF model successfully exhibited diastolic dysfunction, impaired exercise capacity and cardiac hypertrophy with elevated circulating ketone body concentration. Myocardial metabolomics and snRNA-seq identified a profound metabolic shift characterized by the accumulation of long-chain fatty acids and Krebs cycle intermediates, coupled with the transcriptional downregulation of insulin signaling and fatty acid degradation pathways. Although circulating ketone body level was upregulated, *Hmgcs2^ΔiCM^* mice showed no exacerbation of the HFpEF phenotype. In contrast, *Hmgcs2^ΔHep^* mice exhibited significantly aggravated cardiac hypertrophy (HW/TL; *Hmgcs2^flox^*: 7.41 ± 0.87: *Hmgcs2^ΔHep^*: 8.29 ± 0.73; p = 0.0154). Mechanistically, hepatic ketogenesis was required to maintain circulating beta-hydroxybutyrate (BHB) levels, which directly modulated cardiomyocyte metabolism. BHB acted as a metabolic signal to dampen fatty acid overload and facilitate glucose utilization.

**Conclusions:** Our study identifies a critical "liver-heart axis" where hepatic ketogenesis serves as an essential regulator of myocardial metabolic resilience. Impaired hepatic ketogenesis creates a metabolic mismatch that drives pathological cardiac remodeling. These findings highlight the liver as a therapeutic target within the CKM syndrome framework, suggesting that restoring the hepato-cardiac metabolic bridge may ameliorate HFpEF progression.

**What is New?:**

- This study identifies a novel liver–adipose–heart axis that governs myocardial metabolic resilience during the development of heart failure with preserved ejection fraction (HFpEF).
- We demonstrate that while both the liver and heart upregulate ketogenesis under metabolic stress, only hepatic ketogenesis—and not cardiac-intrinsic ketogenesis—is essential for mitigating pathological cardiac remodeling.
- Mechanistically, liver-derived *β* -hydroxybutyrate acts as a critical
- endocrine signal that dampens fatty acid oxidation and facilitates myocardial glucose utilization.

**What Are the Clinical Implications?:**

- Our findings highlight the liver as a central therapeutic target within the cardiovascular–kidney–metabolic (CKM) syndrome framework, where hepatic metabolic failure directly drives cardiac dysfunction.
- Restoring the hepato-cardiac metabolic bridge, through either hepatic-targeted therapies or ketone body supplementation, represents a promising strategy to enhance myocardial metabolic flexibility and ameliorate HfpEF in patients with multi-organ metabolic disorders.

## Introduction

The relationship between metabolic disorders and cardiovascular diseases has drawn increasing attention in recent years. Emerging frameworks such as the cardiovascular–kidney–metabolic (CKM) syndrome underscore the tight bidirectional links among dysglycemia, adiposity, hypertension, and end-organ injury in the heart and vasculature ^1–3^. Heart failure with preserved ejection fraction (HFpEF) and pre-existing cardiac hypertrophy frequently arise in the context of multisystem metabolic diseases—including hypertension, obesity, metabolic dysfunction-associated steatotic liver disease (MASLD), and diabetes—and are shaped by their systemic inflammatory and endocrine milieu ^4–7^. Because the prevalence of these conditions increases with age, and population aging is accelerating globally, clarifying how metabolic abnormalities drive cardiac remodeling and dysfunction has become indispensable^8–11^.

Ketone body metabolism has long been recognized as a key energy-producing system during fasting^12,13^. In mammals, the canonical ketone bodies—acetone, acetoacetate (AcAc) and β-hydroxybutyrate (BHB)—are produced primarily in the liver and transported to peripheral tissues for oxidation. Recent studies have revealed that ketone body metabolism extends beyond mere substrate supply, encompassing diverse signaling and regulatory functions^14,15^. These functions include GPCR-mediated signaling and epigenetic regulation via histone deacetylase (HDAC) inhibition^16,17^. Moreover, ketogenesis is enhanced not only during classical starvation but also in metabolic transition states - such as the neonatal period, weaning, and in certain pathological settings - when a shift between glucose and fatty acid metabolism imposes cellular stress^18^. Specifically, in neonatal liver, ketone body synthesis mitigates excessive fatty acid load and preserves mitochondrial integrity^19^.

The adult heart primarily relies on fatty acid oxidation for ATP generation, but during the development of cardiac hypertrophy and heart failure the metabolic program is remodeled, with reduced fatty acid oxidation and a compensatory increase in carbohydrate use (glycolysis and glucose oxidation)^20,21^. This metabolic remodeling, or metabolic reprogramming, has emerged as a therapeutic target^22,23^. Concurrently, human and animal studies indicate that in heart failure with reduced ejection fraction (HFrEF), circulating ketone levels and myocardial ketone utilization are increased, consistent with recruitment of ketones as a supplemental fuel^24–26^. These observations have spurred interest in exogenous ketone strategies as potential therapies in cardiovascular disease^27^. Notably, recent human flux studies confirm that while fatty acids remain a major contributor to cardiac ATP, the relative contribution of ketones rises in heart failure, reinforcing ketone metabolism as a node of interest for intervention.

However, the molecular mechanisms linking ketone body metabolism to HFpEF and its predisposing condition, cardiac hypertrophy, remain incompletely understood. While it is well established that ketone body utilization is increased in patients with HFrEF, whether a similar alteration occurs in HFpEF is still uncertain and appears to depend on disease phenotype and metabolic context^26,28–30^. Moreover, although exogenous ketone supplementation has been reported to improve cardiac function partly through post-translational modifications and redox signaling, the physiological roles of endogenous ketone production—particularly extra-hepatic ketogenesis—remain largely unexplored in the heart^31–33^. In cardiomyocytes, myocardial ketone synthesis has been implicated in promoting tissue repair and regeneration following ischemic injury^34^. However, whether hepatic or extra-hepatic ketogenesis contributes to the pathogenesis of cardiac hypertrophy or HFpEF, and how this metabolic reprogramming shapes cardiac remodeling, remains unknown. Therefore, in this study, we investigated how metabolic dysfunction triggers cardiac hypertrophy and examined the impact of ketone body metabolism—both systemic and cardiac—on this process.

In this study, we employed an HFpEF model induced by obesity and hypertension to investigate the metabolic crosstalk underlying cardiac remodeling^35^. We found that ketone body synthesis was upregulated in both the heart and the liver during the development of cardiac hypertrophy. Notably, hepatic ketogenesis played a pivotal role in driving the progression of hypertrophy, and the ketone bodies synthesized in the liver modulated circulating fatty acid levels. Furthermore, β-hydroxybutyrate directly influenced cardiomyocytes, enhancing glucose utilization and facilitating metabolic remodeling toward glycolysis. These findings suggest that hepatic ketogenesis mitigates fatty acid overload by coordinating metabolic responses across the liver, adipose tissue, and heart. This inter-organ network mediated by ketone bodies establishes a mechanism of metabolic resilience in cardiomyocytes under stress.

## Methods

### Experimental animals

C57BL/6J background male mice were used. *Hmgcs2^flox^* mice were generated in the previous report^19^. *Alb-cre* mice were provided by Yoshifumi Sato and Kazuya Yamagata. All mice were housed at 23±2 °C, with a relative humidity of 50%–60%, and under a 12-h light: 12-h dark cycle. Genotypes were determined by PCR on tail/fingertip-derived DNA using specific primers. All procedures were performed in accordance with the Kumamoto University animal care guidelines (approval reference no. 2024-064), which conform to the US National Institutes of Health Guide for the Care and Use of Laboratory Animals (publication no. 85-23, revised 1996).

### Echocardiography

Echocardiographic assessment was conducted under isoflurane anesthesia. Transthoracic echocardiography (Toshiba, Japan) was utilized to evaluate cardiac morphology. M-mode measurements included left ventricular (LV) wall thickness and LV systolic and diastolic dimensions. Subsequent calculations yielded LV ejection fraction (EF).

### Blood pressure measurement

Systolic blood pressure was measured in conscious mice using a non-invasive tail-cuff system (MK-2000ST, Muromachi kikai, Japan). To minimize stress artifacts, mice were acclimatized to the procedure prior to data collection. The mean of at least five consecutive measurements was calculated for analysis

### Hemodynamic test

Each mouse was anesthetized with 2 % isoflurane and endotracheally intubated to maintain mechanical ventilation. After performing a midline sternotomy to expose the heart, a 1.4-Fr pressure–volume catheter (SPR-839; Millar Instruments, USA) was inserted into the left ventricle via the apex for pressure–volume (PV) loop measurements using a Millar MPVS Ultra Pressure–Volume System. Baseline PV loops were recorded under steady-state conditions. Subsequently, transient occlusion of the inferior vena cava was performed to obtain preload-independent PV relationships.

### Treadmill exercise test

After 3 days of acclimatization to the treadmill for 3 days prior to testing, treadmill exercising tests were conducted using a five-lane rodent treadmill (LE8710MRTS+, Panlab, Spain) set at a 20° incline. On the test day, running speed was increased by 2 m/s every 2 min, starting from 14 m/min, until the mouse reached exhaustion, defined as the inability to continue running for >10 s despite mild electrical stimulation (0.2 mA). Total running distance and time were recorded using SeDaCom software (Panlab).

### Histological analysis

Hematoxylin-eosin (HE) staining was performed using paraffin-embedded sections (10 µm)(K.I. Stainer Inc., Kumamoto, Japan). For, immunohistochemical analysis was performed on paraffin-embedded sections (10 µm). The antibodies used were as follows: 4’,6-Diamidino-2-phenylindole, dihydrochloride, solution (DAPI: Dojindo, D523), Alexa Fluor 488-conjugated wheat germ agglutinin (WGA: ThermoFisher, W11261), Isolectin GS-IB4 From Griffonia simplicifolia, Alexa Fluor™ 647 Conjugate (IB4: ThemoFisher, I32450) and Phalloidin-TRITC (Sigma Aldrich, P1951).

### Western blot analysis

Western blots were performed with an SDS-PAGE system. Blotted gels (Bio-Rad, TGX Stain-Free™ FastCast™ Acrylamide Solutions) were transferred to PVDF membranes using the Trans-Blot Turbo Transfer System (Bio-Rad). Membranes were immersed in 5% skim-milk containing blocking solution and reacted with a specific primary antibody (Hexokinase I (C35C4) Rabbit mAb: CST, #2024).

### Measurement of beta-hydroxybutyrate (βOHB)

For whole blood and serum monitoring, we used FreeStyle Precision Neo and FreeStyle Precision Blood β-ketone test strips (Abbott). Whole blood samples were obtained by tail cutting or by decapitation. For serum assessment, whole blood samples were collected in Capillary Blood Collection Tubes (Terumo) and were centrifuged at 3,000 g at room temperature (RT) for 10 min.

### Nuclei isolation

Frozen tissue samples (20–40 mg) were sectioned into fine pieces using a razor blade and immediately transferred to a chilled glass homogenization vessel. Mechanical dissociation was performed using a Potter–Elvehjem homogenizer with the pestle submerged beneath the buffer surface to minimize air exposure. Homogenization was conducted with 5 vertical strokes at 1,000 rpm under ice-cold conditions. The resulting homogenate was mixed with sucrose buffer (composition described below) and adjusted to a final sucrose concentration of 27% (w/v). The sample was then carefully loaded into an ultracentrifuge tube to generate a discontinuous Percoll (MP Biomedicals, 592-09051) density gradient consisting of, from top to bottom, 12%, 27% (sample layer), 31%, and 35% (w/v) sucrose fractions. Gradient separation was performed using an ultracentrifuge (e.g., Beckman Coulter Optima MAX-TL) at 24,000 rpm for 10 min at 4°C. After centrifugation, intact nuclei were recovered from the densest interface/bottom fraction. The purified nuclear suspension was collected using a chilled glass pipette, transferred to a low-binding tube, and processed for downstream applications including transcriptomic analysis.

### Flow cytometry and nuclei sorting

Nuclei sorting and flow cytometric analyses were performed using the high-speed cell sorter MoFlo XDP operated in accordance with the manufacturer’s specifications. Purified nuclei were incubated with fluorescence-conjugated antibodies targeting the nuclear envelope and cardiomyocyte-specific markers: AF488-conjugated anti-Lamin A/C antibody (CST, #8617) and AF647-conjugated anti-PCM1 antibody. Antibody staining was conducted in 1% BSA/PBS for 30 min at 4°C. Fluorescence signal intensities were measured using a calibrated flow cytometer equipped with 488-nm and 640-nm excitation lasers. Forward and side scatter parameters were used to gate intact singlet nuclei while excluding debris and doublets. FCS files were analyzed using standard cytometry software, and nuclei positive for both Lamin A/C and PCM1 were isolated for downstream molecular assays.

### RamDA sequencing analysis

Nuclei-adapted transcriptomic profiling was performed based on a modified RamDA-seq protocol following the strategy reported by Hayashi et al.^36^ Briefly, 1,000 sorted nuclei were collected into low-binding tubes containing lysis buffer supplemented with RNase inhibitor and immediately processed through the nuclei-optimized RamDA reaction. Sequencing libraries were constructed using the NEBNext Ultra II DNA Library Prep Kit (NEB, #E7805), with index incorporation and amplification performed according to the manufacturer’s guidelines. Libraries were sequenced on an Illumina platform using 150-bp paired-end chemistry on the high-throughput sequencer NovaSeq 6000. Raw FASTQ files were processed using established quality control and preprocessing tools. Adapter sequences and low-quality bases were removed using gradient-aware trimming software (e.g., Trimmomatic). Read quality metrics before and after trimming were assessed using the NGS QC program FastQC. Transcript-level quantification was performed using the alignment-free estimator Salmon to obtain gene-level count matrices against the appropriate species transcriptome reference. Differential expression analysis was completed using the RNA-seq statistical framework DESeq2. Pathway-level enrichment was evaluated using the gene-set testing method Gene Set Enrichment Analysis implemented through either fgsea or the standalone GSEA toolset (method name not repeated here). All analyses were conducted in the R statistical environment (script details available upon request).

### GC-MS analysis

A frozen tissue or liquid sample was plunged into Milli-Q water/methanol/chloroform (1:2.5:1) containing internal standards (isopropylmalic acid). The sample was homogenized at 2,300 rpm for 30 s with a bead beater-type homogenizer (Beads crusher µT-12, Titec Corporation). The homogenized sample was shaken at 1,200 rpm and 37 °C for 30 min and centrifuged at 16,000 g and 4 °C for 3 min. Subsequently, 225 µL of the upper aqueous layer was transferred to a new tube. The remaining sample was mixed with 200 µL Milli-Q water and centrifuged again at 16,000 g and 4°C for 3 min. The upper water layer was mixed with the aqueous layer. The solution was evaporated using a centrifugal evaporator (DNA120OP230, ThermoFisher Scientific) and lyophilized (FD-1-84A, FST). The sample was derivatized with methoxamine/pyridine and N-methyl-N-trimethylsilyltrifluoroacetamide and analyzed on a Shimadzu TQ8050 GC–MS/MS (Shimadzu, Kyoto, Japan). The chromatograms and mass spectra were analyzed utilizing the GC–MS solution software 4.50 (Shimadzu, Kyoto, Japan). Compounds were determined with the Smart Metabolites Database Ver.2 (Shimadzu, Kyoto, Japan).

### Cell culture and In Vitro hypertrophy model

H9C2, rat cardiomyocyte-derived cells, were maintained in Dulbecco’s Modified Eagle Medium (DMEM) supplemented with 10% fetal bovine serum (FBS) and 1% penicillin/streptomycin at 37℃ in a humidified atmosphere containing 5% CO2. To induce cardiomyocyte hypertrophy, H9C2 cells were first serum-starved for 24 hours in DMEM (Serum-free medium) and the cells were treated with 100 μM Phenylephrine ((R)-(-)-Phenylephrine Hydrochloride, FUJIFILM Wako Pure Chemical Corporation, Japan) for 48 hours. Control cells were treated with an equal volume of PBS.

### Measurement of Oxygen Consumption Rate (OCR)

H9C2 cell line was used for the analysis. Measurement of oxygen consumption rate was described previously 7. XF HS mini Extracellular Flux Analyzer (Seahorse Bioscience) was used for the analysis. During the real-time measurement, inhibitors of respiratory chain components, oligomycin (1 μg/ml, the complex V inhibitor), carbonyl cyanide-p-trifluoromethoxyphenylhydrazone (FCCP) (1 μM, respiratory uncoupler), rotenone and antimycin A (100nM and 10 μM, the complex I and III inhibitors) were administered sequentially. OCR for Basal respiration minus proton leak (rate prior to FCCP injection minus non-mitochondrial respiration) was determined as ATP consumption. Maximal respiration was determined by rate during FCCP administration minus rate after rotenone/antimycin-added states. When fatty acid oxidation was assessed, we mixed 150 μM palmitate-BSA or BSA during the real-time measurement of OCR.

### Statistical analysis

Data for normally distributed continuous variables are expressed as means ± SD. Two-group comparisons were carried out with Welch’s t-test. Statistical analyses were performed with Prism 7 (GraphPad Software Inc). Two-way ANOVA followed by Sidak’s multiple comparisons test was used for factor analysis.

## Result

### Elevation of Free Fatty Acids and Ketone Bodies in the HFpEF Model

As previously reported, the murine model with coexisting obesity and hypertension drives to heart failure with preserved ejection fraction (HFpEF)^35^. We assessed the impact of ketone body metabolism using this HFpEF model. Adult wild-type mice (8–10 weeks old) were subjected to combined stress—defined as a high-fat diet and water containing 0.5 g/L NG-Nitro-L-arginine methyl ester hydrochloride (L-NAME)—for 15 weeks (Figure 1a). This intervention significantly increased body weight (Control: 27.1 ± 1.6 g, n=10; HFpEF: 40.9 ± 6.5 g, n=10, p < 0.0001) and systolic blood pressure (Control: 111.6 ± 13.5 mmHg; HFpEF: 135.3 ± 21.8 mmHg, p = 0.0106) (Figure 1b and c). An increase in blood glucose levels was also observed in the HFpEF model, with a significant elevation particularly under *ad libitum* feeding conditions (*Ad lib.*) (Figure 1d).

**Figure 1.**
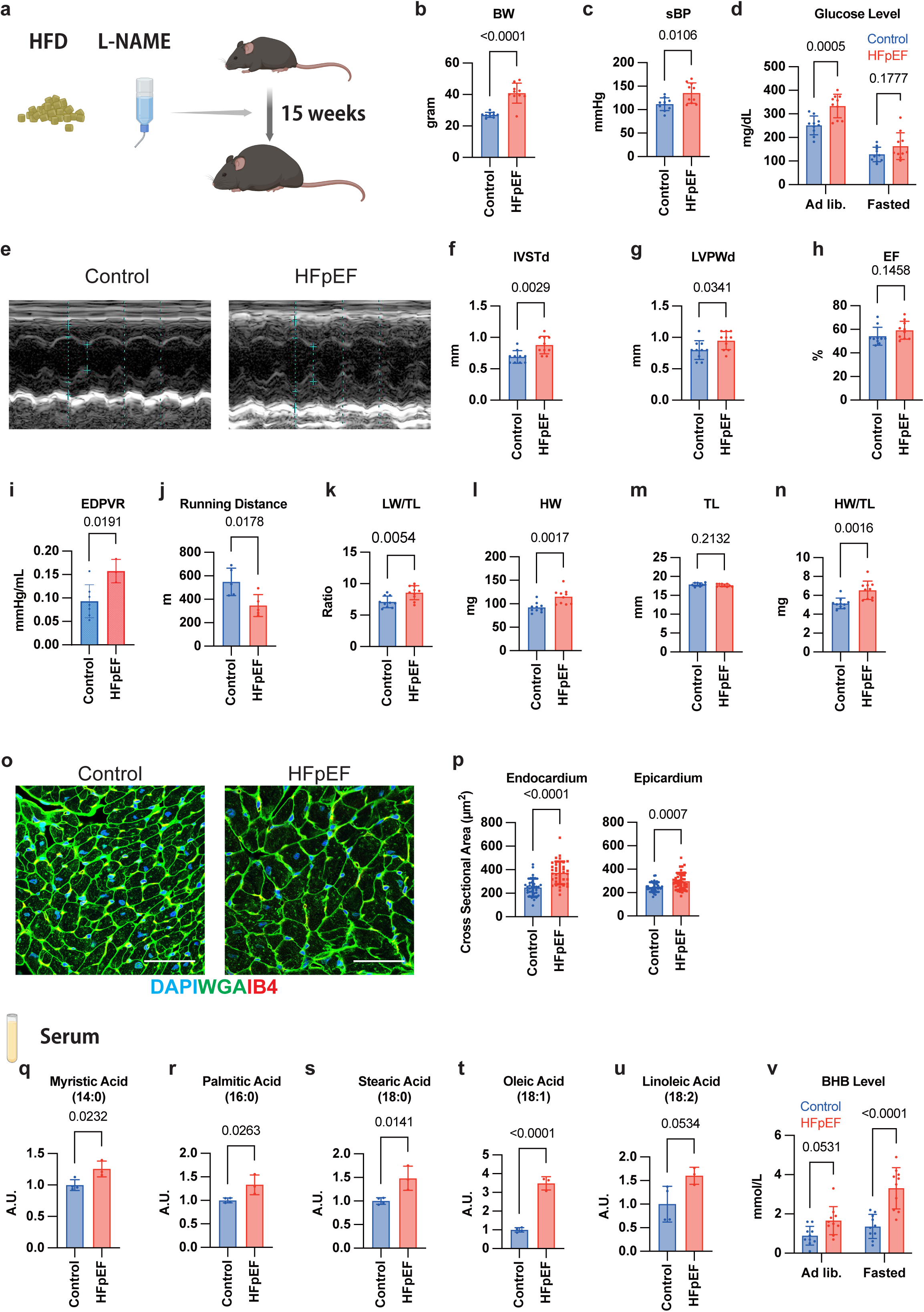
The HFpEF model enhances ketogenesis. a, A schematic showing the combined stress model. b, body weight. c, systolic blood pressure (sBP). D, Blood glucose level. E, Representative results of echocardiography. f, interventricular septum thickness at diastole (IVSTd). g, left ventricular posterior wall thickness at diastole (LVPWd). h, ejection fraction (EF). i, end-diastolic pressure volume relationship (EDPVR) measured by Pressure-Volume loop system catheter. j, symptom-limited running distance measured by the treadmill test k, Lung weight l, heart weight (HW). m, tibial length (TL). n, adjusted heart weight by tibial length (HW/TL). o, heart immunohistochemical staining with 4’,6-diamidino-2-phenylindole (DAPI), wheat germ agglutinin lectin (WGA), and isolectin B4 (IB4). Scale bar: 50 μm. p, cross sectional areas of endocardial cardiomyocytes and epicardial cardiomyocytes. q-u, relative serum concentration of fatty acid. q, myristic acid. r, palmitic acid. s, stearic acid. t, oleic acid. u, linoleic acid. v, relative serum concentration of beta hydroxyburyrate under ad-libitum (Ad lib.) or fasted feeding. Welch’s t-tests and two-way ANOVA analysis were used for statistical analysis.

Echocardiography revealed that the HFpEF condition increased the interventricular septum thickness at diastole (IVSTd; Control: 0.69 ± 0.10 mm, n=10; HFpEF: 0.88 ± 0.14 mm, n=10, p = 0.0029) and left ventricular posterior wall thickness at diastole (LVPWd; Control: 0.80 ± 0.15 mm; HFpEF: 0.95 ± 0.14 mm, p = 0.0341) with preserved ejection fraction (EF; Control: 54.1 ± 7.6 %; HFpEF: 59.3 ± 7.6 %, p = 0.146) (Figure 1e-h). Pressure-volume loop analysis also demonstrated a significant increase in the end-diastolic pressure-volume relationship (EDPVR) in the HFpEF group (Control: 0.093 ± 0.035 mmHg/μL, n=7 vs. HFpEF: 0.158 ± 0.025 mmHg/μL, n=4; p = 0.0191) (Figure 1i). A treadmill-based exhaustion test revealed a reduced exercise capacity in the HFpEF model (Control: 548.6 ± 117.2, n=5 vs. HFpEF: 346.8 ± 93.6, n=5; p = 0.0178) (Figure 1j). The lung weight to tibial length ratio, an indicator of pulmonary congestion, was significantly increased (Control: 7.11 ± 0.91, n=10 vs. HFpEF: 8.56 ± 1.12, n=10; p = 0.0054) (Figure 1k). Heart weight and the heart weight to tibial length ratio showed a significant increase in the HFpEF group (HW: Control: 92.0 ± 10.5 mg vs. HFpEF: 115.0 ± 16.0 mg; p = 0.0017; HW/TL: Control: 5.15 ± 0.56 vs. HFpEF: 6.53 ± 0.97; p = 0.0016) (Figure 1l-n). Histological analysis revealed increased cardiomyocyte cell size in the HFpEF group (Figure 1n). Quantification analysis confirmed the increased cardiomyocyte cell size both in endocardium (Control: 249.1 ± 76.3 μm^2^: HFpEF: 373.8 ± 96.2 μm^2^; p < 0.0001, Figure 1o) and epicardium (Control: 248.2 ± 41.5 μm^2^: HFpEF: 296.7 ± 71.9 μm^2^; p = 0.0007 Figure 1p). From these data, we confirm that the HFpEF model induces significant cardiac hypertrophy and meets the criteria for a heart failure model. To investigate alterations in the lipid profile associated with the HFpEF model, we performed metabolomic analysis to measure circulating free fatty acids. This analysis revealed that multiple long-chain fatty acids, both saturated and unsaturated, were elevated in the HFpEF model (Figure 1q-u). In addition, an increase in circulating ketone bodies was observed, with serum β-hydroxybutyrate (β-OHB) levels significantly elevated in the HFpEF model, particularly under fasting conditions (Figure 1v).

### Metabolic changes in the heart with HFpEF model

In the subsequent analysis, we examined metabolic alterations within the hearts of HFpEF model mice. β-Hydroxybutyrate levels were markedly elevated in cardiac tissue, whereas a decreasing trend was observed for glycolytic intermediates, including glucose and glucose-6-phosphate. In contrast, Krebs cycle intermediates—such as citrate, isocitrate, and malate—were increased (Figure 2a). These findings suggested enhanced acetyl-CoA generation via lipolysis. Therefore, we next assessed the levels of free fatty acids in cardiac tissue. Metabolomic profiling revealed elevations in both saturated and unsaturated free fatty acids, mirroring the pattern observed in the circulation of the HFpEF model (Figure 2b–g).

**Figure 2.**
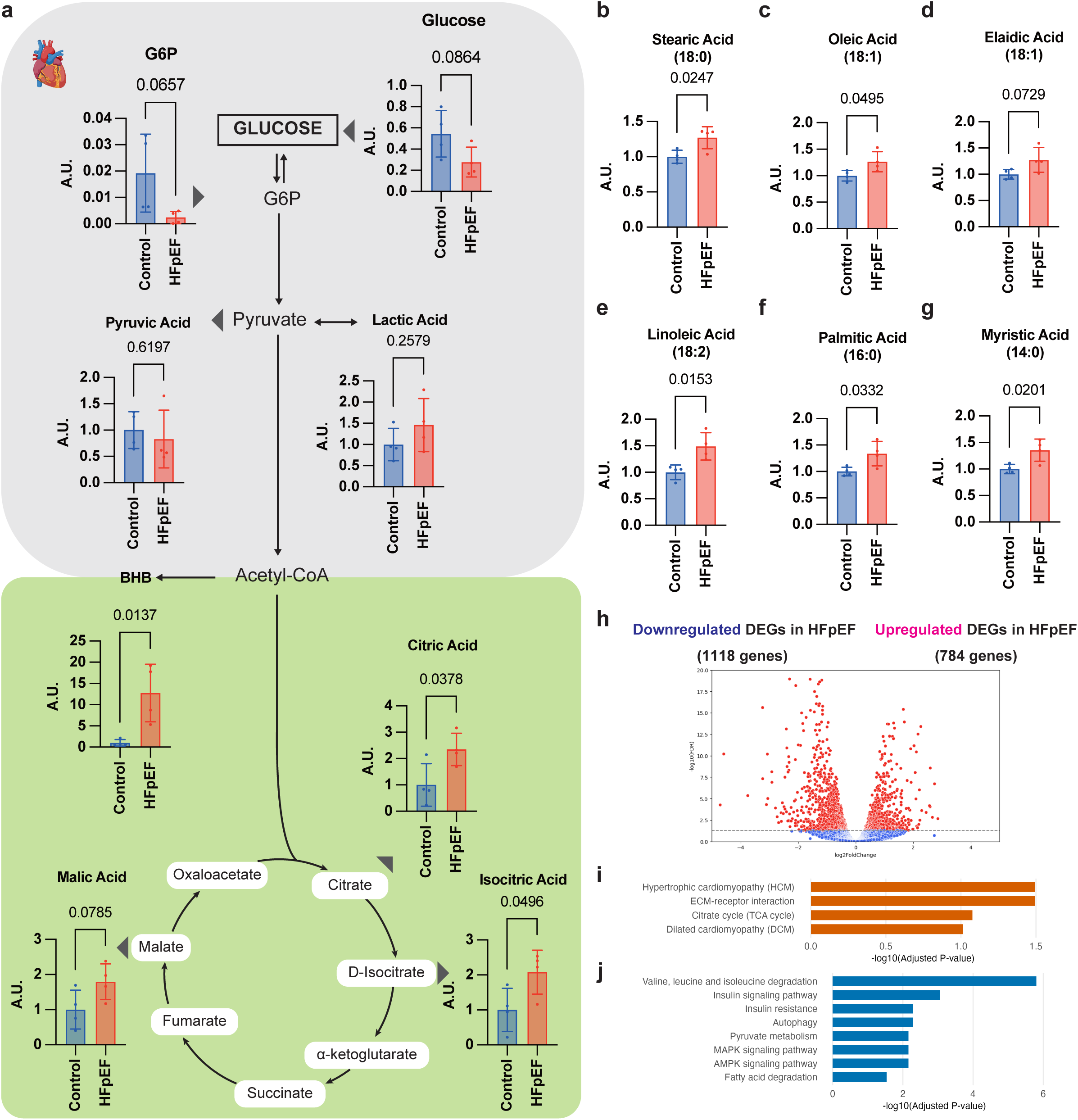
Metabolic and Transcriptional Landscape of the HFpEF Heart. a, A schematic showing the metabolic pathway and concentration of each metabolite including glucose-6-phosphate(G6P), glucose, pyruvic acid, lactic acid, β-hydroxybutyrate (BHB), citric acid, isocitric acid and malic acid. b-g, relative concentration of fatty acid in heart tissue. h, Volcano plot showing differentially expressed genes (DEGs) between HFpEF and control hearts. The x-axis represents the log2 fold change (log2FC), and the y-axis represents the negative log10 adjusted P-value (-log10(Adjusted P-value)). Red dots indicate significantly altered genes in HFpEF. i-j, Bar plots displaying top enriched KEGG pathways from Gene Set Enrichment Analysis (GSEA) of DEGs in HFpEF. Enriched pathways for upregulated (i) and downregulated (j) DEGs in HFpEF. Welch’s t-tests were used for statistical analysis.

Cardiomyocyte-specific RNA sequencing was performed to capture comprehensive transcriptional changes, with a particular focus on genes related to metabolic enzymes. Differential gene expression (DEG) analysis revealed significant transcriptional alterations, identifying 1,118 downregulated and 784 upregulated transcripts (Figure 2h). Subsequently, pathway analysis demonstrated that the HFpEF model exhibited an enrichment of pathways associated with cardiomyopathy and a clear increase in TCA cycle-related processes (Figure 2i). Conversely, the significantly downregulated pathways included those critical for insulin signaling and fatty acid degradation (Figure 2j).

Collectively, these findings demonstrate that the HFpEF model is characterized by a reduction in myocardial glycolytic intermediates but a concurrent accumulation of TCA cycle intermediates and fatty acids. Crucially, this metabolic shift is further supported at the transcriptional level by the downregulation of fatty acid degradation pathways.

### Loss of intrinsic cardiac ketogenesis does not exacerbate cardiac hypertrophy

To elucidate the impact of ketone body metabolism on hypertrophic pathogenesis, we first examined the role of cardiac ketogenesis. We generated cardiomyocyte-specific, inducible *Hmgcs2* knock out mice (hereafter referred to as *Hmgcs2^ΔiCM^*) by crossing *Hmgcs2^flox/flox^* mice (*Hmgcs2^flox^*) with MHC-merCremer mice. To induce Cre recombination, *Hmgcs2^flox^* and *MHC-Hmgcs2^ΔiCM^* mice were administrated tamoxifen at a dosage of 20 mg/kg for 5 consecutive days (Figure 3a). We confirmed the successful and specific deletion of *Hmgcs2* in the heart. Western blot analysis demonstrated a significant reduction of Hmgcs2 protein in the heart of *Hmgcs2^ΔiCM^* mice compared to *Hmgcs2^flox^* control mice (Figure 3b). Crucially, Hmgcs2 protein level in the liver remained unchanged between the two genotypes (Figure 3c), confirming the tissue specificity of the gene deletion.

**Figure 3.**
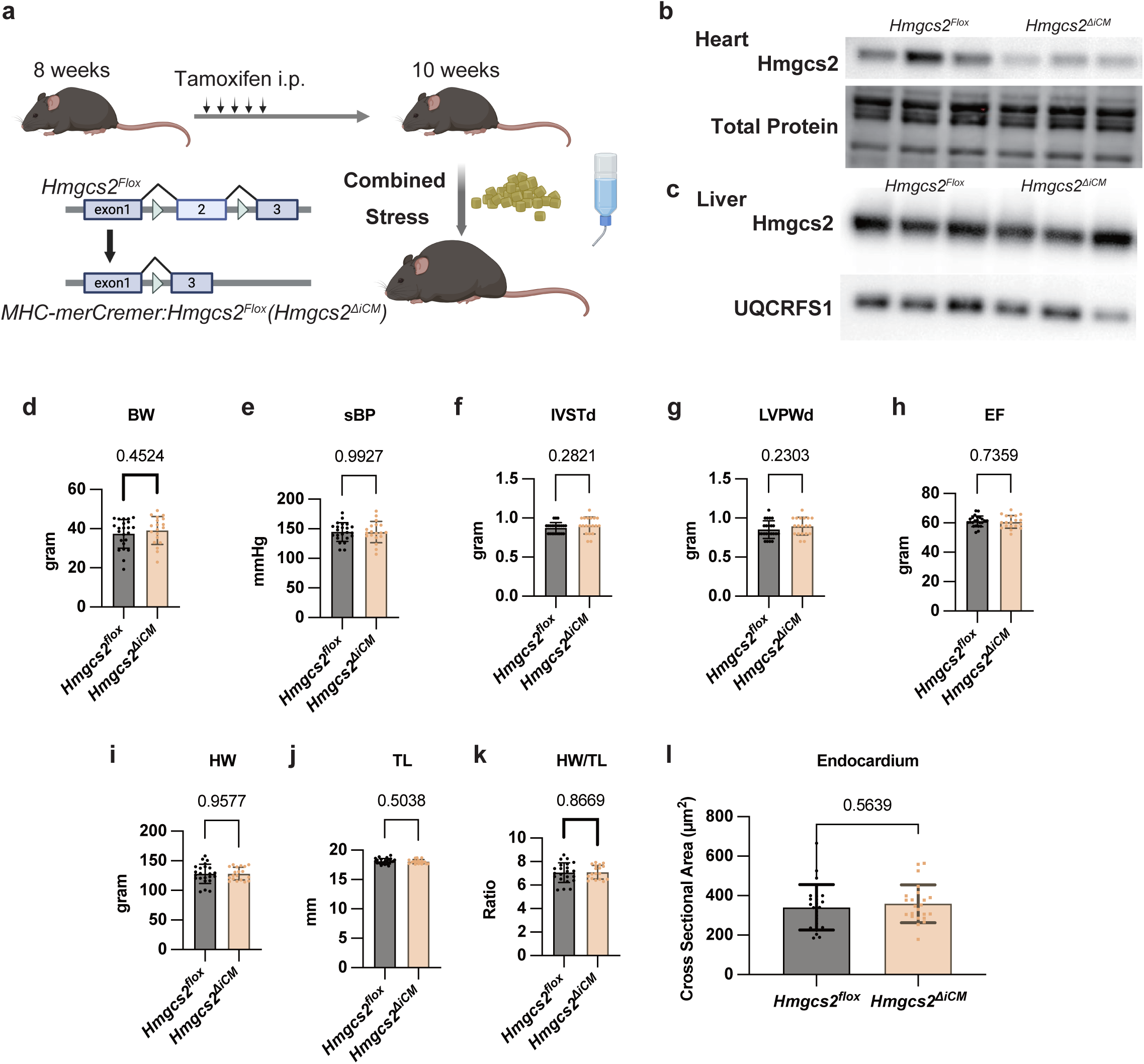
Loss of intrinsic cardiac ketogenesis does not exacerbate cardiac hypertrophy. a, A schematic showing the deletion of Hmgcs2 in cardiomyocytes and induction of the combined stress. b, the image of western blotting analysis of heart tissues with HMG-CoA synthase2 (Hmgcs2) and total protein. c, the image of western blotting analysis of liver tissues with HMG-CoA synthase2 (Hmgcs2) and ubiquinol-cytochrome c reductase, Rieske iron-sulfur polypeptide 1 (UQCRFS1). d, body weight (BW). e, systolic blood pressure (sBP). f, interventricular septum thickness at diastole (IVSTd). g, left ventricular posterior wall thickness at diastole (LVPWd). h, ejection fraction (EF). i, heart weight (HW). j, tibial length (TL). k, adjusted heart weight by tibial length (HW/TL). l, cross sectional areas of endocardial cardiomyocytes. Welch’s *t*-tests were used for statistical analysis.

Following the successful deletion of *Hmgcs2* in the myocardium, we subjected both *Hmgcs2^ΔiCM^* and *Hmgcs2^flox^* mice to combined stress. After 15 weeks of combined stress, no significant change were observed in body weight (BW) and systolic blood pressure(sBP) between the two genotypes (BW; *Hmgcs2^flox^*: 37.4 ± 7.4 g, n=23; *Hmgcs2^ΔiCM^*: 39.1 ± 7.1 g, n=19, p = 0.452, sBP; *Hmgcs2^flox^*: 144.5 ± 15.7 mmHg, n=23; *Hmgcs2^ΔiCM^*: 144.5 ± 17.9 mmHg, n=19, p = 0.993) (Figure 3d and e). Furthermore, detailed echocardiography analysis revealed that the deletion of *Hmgcs2* did not affect the structural or functional parameter (IVSTd; *Hmgcs2^flox^*: 0.87 ± 0.69 mm, n=23; *Hmgcs2^ΔiCM^*: 0.91 ± 0.11 mm, n=19, p = 0.282, LVPWd; *Hmgcs2^flox^*: 0.85 ± 0.11 mm, n=10; *Hmgcs2^ΔiCM^*: 0.89 ± 0.11 mm, n=19, p = 0.230, EF; *Hmgcs2^flox^*: 61.1 ± 3.6 %, n=23; *Hmgcs2^ΔiCM^*: 60.6 ± 4.3 %, n=19, p = 0.736) (Figure 3f, g, and h). Consistent with the echocardiography findings, heart weight(HW) was comparable between the control and knockout groups (HW; *Hmgcs2^flox^*: 128.0 ± 16.3 mg, n=23: *Hmgcs2^ΔiCM^*: 128.2 ± 10.8 mg, n=19; p = 0.958, HW/TL; *Hmgcs2^flox^*: 7.06 ± 0.85, n=22: *Hmgcs2^ΔiCM^*: 7.09 ± 0.59, n=18; p = 0.867) (Figure 3i, j, and k). Furthermore, histological analysis showed that cardiomyocyte cell size was indistinguishable between *Hmgcs2^flox^* and *Hmgcs2^ΔiCM^* mice (Endocardial cardiomyocyte cell size; *Hmgcs2^flox^*: 340.7 ± 114.9 μm^2^: *Hmgcs2^ΔiCM^*: 358.9 ± 96.1 μm^2^; p = 0.564) (Figure 3l).

Taken together, these data strongly indicate that the cardiomyocyte-specific ketogenesis is not a major determinant of structural hypertrophy under the tested chronic stress condition.

### Loss of hepatic ketogenesis exacerbates cardiomyocyte hypertrophy

Next, we assessed the impact of hepatic ketogenesis, the major source of circulating ketone bodies, on cardiac hypertrophy. To specifically investigate the role of Hmgcs2 in the liver, we generated hepatocyte-specific *Hmgcs2* knockout mice (hereafter referd to as *Hmgcs2^ΔHep^*) by crossing *Alb-cre mice and Hmgcs2^flox/flox^* (*Hmgcs2^flox^*). We first confirmed the successful and specific gene deletion in the liver (Figure 4a and 4b). Crucially, serum ketone body concentration was significantly reduced in *Hmgcs2^ΔHep^* (β-hydroxybutyrate; *Hmgcs2^flox^*: 5.34 ± 0.75 mmol/L: *Hmgcs2^ΔHep^*: 0.34 ± 0.11 mmol/L; p < 0.0001) (Figure 4c). These results confirmed that *Hmgcs2^ΔHep^* mice serve as an effective model for examining the effects of impaired hepatic ketogenesis *in vivo*.

**Figure 4.**
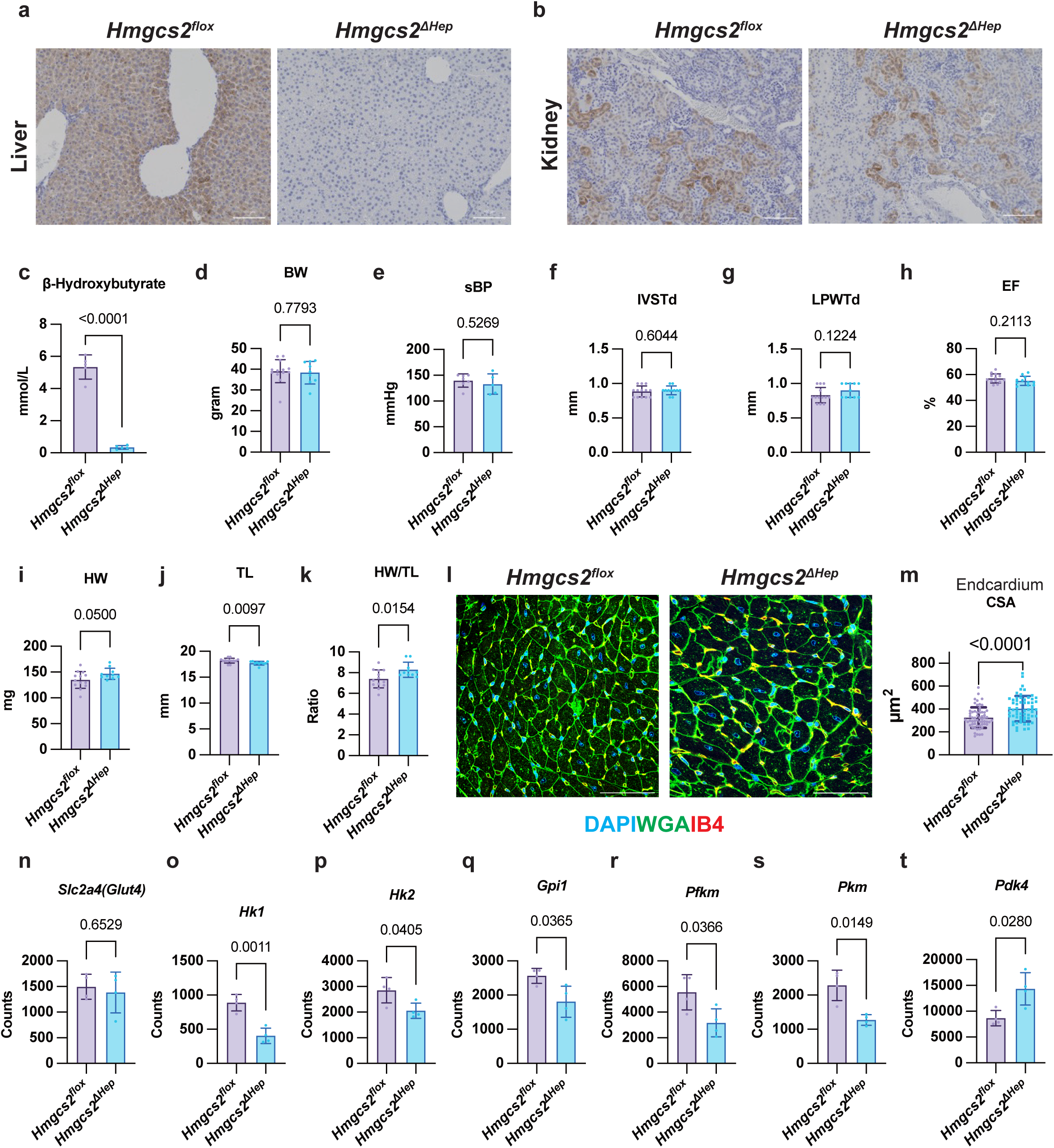
Loss of hepatic ketogenesis exacerbates cardiac hypertrophy. a, Immunohistochemical image of livers with HMG-CoA synthase 2 (Hmgcs2). b, Immunohistochemical image of kidneys with Hmgcs2. c, Concentration of β-hydroxybutyrate in serum. d, body weight (BW). e, systolic blood pressure (sBP). f, interventricular septum thickness at diastole (IVSTd). g, left ventricular posterior wall thickness at diastole (LVPWd). h, ejection fraction (EF). i, heart weight (HW). j, tibial length (TL). k, adjusted heart weight by tibial length (HW/TL). l, heart immunohistochemical staining with 4’,6-diamidino-2-phenylindole (DAPI), wheat germ agglutinin lectin (WGA), and isolectin B4 (IB4). Scale bar: 50 μm. m, cross sectional areas of endocardial cardiomyocytes. n-t, relative expression in cardiac tissue confirmed by RNA sequencing analysis n, Slc2a4, encoding glucose transporter 4 (GLUT4), o, hexokinase1 (Hk1), p, hexikinase2 (Hk2), q, glucose-6-phosphate isomerase (Gpi), r, phosphofruktokinase-muscle (PFKm), s, pruvate kinase m (Pkm) t, pyruvate dehydrogenase lipoamide kinase isozyme 4 (PDK4). Welch’s t-tests were used for statistical analysis.

Subsequently, we subjected the *Hmgcs2^ΔHep^* and *Hmgcs2^flox^* control mice to 15 weeks of combined stress and assessed the cardiac phenotypes. Similar to the control model, body weight and systolic blood pressure did not show the significant changes between *Hmgcs2^flox^* and *Hmgcs2^ΔHep^* (BW; *Hmgcs2^flox^*: 39.1 ± 5.6 g, n=13; *Hmgcs2^ΔHep^*: 38.4 ± 5.5 g, n=9, p = 0.779, sBP; *Hmgcs2^flox^*: 139.7 ± 12.9 mmHg, n= 7; *Hmgcs2^ΔHep^*: 133.0 ± 19.6 mmHg, n=5, p = 0.527) (Figure 4d and e) and echocardiography analysis revealed that preserved ejection fraction with cardiac hypertrophy in both groups (IVSTd; *Hmgcs2^flox^*: 0.89 ± 0.080 mm, n=13; *Hmgcs2^ΔHep^*: 0.90 ± 0.063 mm, n=11, p = 0.604, LVPWd; *Hmgcs2^flox^*: 0.83 ± 0.11 mm, n=13; *Hmgcs2^ΔHep^*: 0.90 ± 0.10 mm, n=11 p = 0.122, EF; *Hmgcs2^flox^*: 57.0 ± 3.6 %, n=13; *Hmgcs2^ΔHep^*: 55.2 ± 3.5 %, n=11, p = 0.211) (Figure 4f, g, and h). However, we observed a significant increase in the indices of cardiac hypertrophy in *Hmgcs2^ΔHep^* mice. Heart weight (HW) exhibited a strong trend toward increase in *Hmgcs2^ΔHep^* mice (HW; *Hmgcs2^flox^*: 134.9 ± 16.3 mg, n=13: *Hmgcs2^ΔHep^*: 146.7 ± 10.9 mg, n=10; p = 0.050). Notably, the heart weight adjusted ratio by tibial length was significantly elevated in *Hmgcs2^ΔHep^* mice (TL; *Hmgcs2^flox^*: 18.2 ± 0.46 mm, n=13: *Hmgcs2^ΔHep^*: 17.7 ± 0.36 mm, n=10; p = 0.0097, HW/TL; *Hmgcs2^flox^*: 7.41 ± 0.87, n=13: *Hmgcs2^ΔHep^*: 8.29 ± 0.73, n=10; p = 0.0154) (Figure 4i, j, and k). Furthermore, histological analysis confirmed a marked exacerbating of cellular hypertrophy in *Hmgcs2^ΔHep^* (Endocardial cardiomyocyte cell size; *Hmgcs2^flox^*: 322.8 ± 88.3 μm^2^: *Hmgcs2^ΔHep^*: 444.0 ± 118.5 μm^2^; p < 0.0001) (Figure 4l, m). Taken together, these data clearly demonstrate that the impairment of hepatic ketogenesis exacerbated the microscopic cardiomyocyte hypertrophy.

Based on the observed exacerbation of hypertrophy in *Hmgcs2^ΔHep^* mice, we hypothesized that ketone bodies influence the metabolic reprogramming that occurs during cardiac hypertrophy. It has been previously reported that myocardial metabolic remodeling—specifically a shift from fatty acid oxidation toward glycolysis—acts as a compensatory and protective mechanism in hypertrophic hearts^7^. To test this hypothesis, we performed RamDA-seq analysis on myocardial tissue. This analysis revealed a significant downregulation of several glycolytic key enzymes in *Hmgcs2^ΔHep^* mice (Figure 4n–s). Conversely, Pyruvate Dehydrogenase Kinase4 (*Pdk4*), a key enzyme involved in metabolic switching between carbohydrate and fatty acid utilization, was upregulated in *Hmgcs2^ΔHep^* (Figure 4t). These results suggest that the defect in hepatic ketogenesis leads to the loss of metabolic shift and dominance of fatty acid metabolism in the hypertrophic myocaridium.

### Hepatocyte destruction and serum fatty acid level increase occur in *Hmgcs2^ΔHep^*

Previous reports have indicated that the failure of ketogenesis is associated with metabolic-associated fatty liver disease (MAFLD)^37–39^. To investigate the causes underlying the persistent fatty acid metabolism-dominant metabolic phenotype observed in the *Hmgcs2^ΔHep^* heart, we proceeded with the analysis of the liver and serum. Upon reevaluation of liver tissue, only mild fat deposition was observed in the *Hmgcs2^ΔHe^****^p^*** under non-stress conditions (Figure 5a). However, combined stress resulted in increased ectopic fat deposition in both groups (Figure 5b). Focusing on liver findings under combined stress, the architecture in the perivenous zone 3, where ketogenesis actively occurred, was relatively preserved in *Hmgcs2^flox^*, whereas *Hmgcs2^ΔHe^****^p^*** showed marked degeneration in this area (Figure 5b, arrowhead).

**Figure 5.**
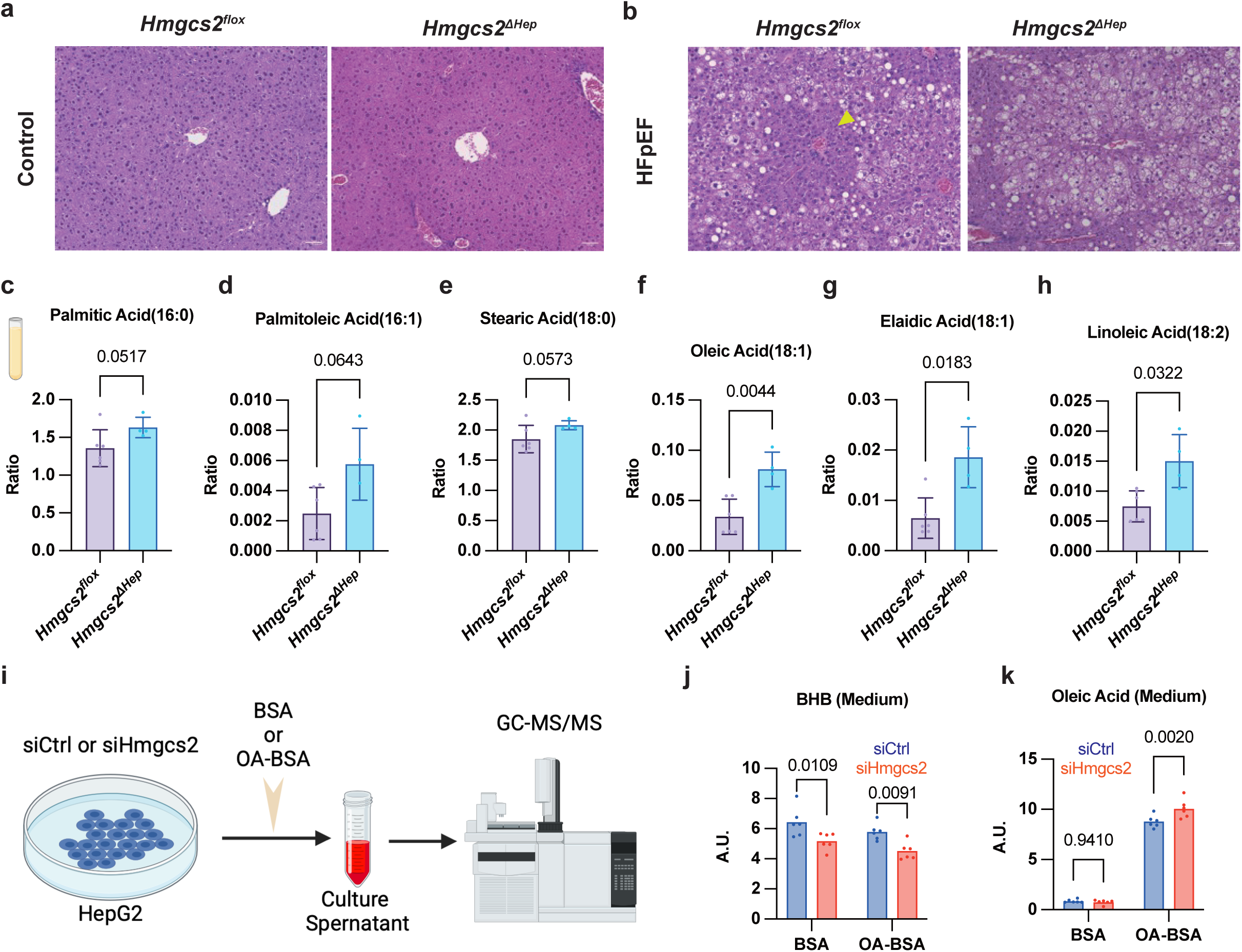
Hepatocyte destruction and serum fatty acid level increase occur in Hmgcs2ΔHep. a, HE staining image of liver under no stress. b, HE staining image of liver under combined stress. c-h, relative serum concentration of fatty acids. I, experimental schema: HepG2 cells transfected with Hmgcs2 or control siRNA, cultured with BSA in the presence or absence of oleic acid, and the supernatant was subjected to metabolome analysis using GC-MS/MS. j, relative concentration of β-hydroxybutyrate in each culture or siRNA. k, relative concentration of oleic acid in each culture or siRNA. Welch’s t-tests and two-way ANOVA analysis were used for statistical analysis.

Furthermore, serum fatty acid analysis revealed a significant increase in the concentration of both saturated and unsaturated fatty acids in *Hmgcs2^ΔHe^****^p^*** serum (Figure 5c-h). These findings suggest that impaired lipid utilization due to insufficient hepatic ketogenesis, coupled with hepatocyte damage (especially in zone 3), leads to an elevated concentration of fatty acids in the blood. This hyperlipidemia is the likely mechanism driving the persistence of the fatty acid metabolism-dominant metabolic phenotype in the heart. These results collectively indicate that a loss of ketogenesis reduces systemic fatty acid metabolic capacity, leading to impaired clearance of fatty acids in the blood. Based on *in vivo* findings, we then performed experiments using HepG2, human hepatoblastoma cells. HepG2 cells were transfected with Hmgcs2 siRNA (siHMGCS2) or Control (siCtrl), and then treated with BSA containing oleic acid or BSA without oleic acid, followed by metabolomics analysis of the culture supernatant by GC-MS/MS (Figure 5i). Consistent with the *in vivo* observations, β-hydroxybutyrate levels significantly decreased in siHMGCS2 cells regardless of the presence of oleic acid in the medium (Figure 5j). Conversely, in the medium supplemented with oleic acid, a higher concentration of residual oleic acid remained in the siHMGCS2 group (Figure 5k), which indicated that the decreased ketogenesis impaired fatty acids metabolism under a condition of abundant fatty acid supply. These *in vitro* results confirmed that hepatic ketogenesis contributes substantially to the utilization of fatty acids in the liver, and this capacity significantly affects blood fatty acid concentration, which might impact cardiac metabolism.

### β-Hydroxybutyrate directly suppresses fatty acid utilization and promotes glucose utilization in cardiomyocytes

Next, we investigated the direct effects of β-hydroxybutyrate (BHB) on cardiomyocyte metabolism to further verify their influence of circulating ketone bodies. First, H9C2 cells were treated with 100μM phenylephrine to induce cardiac hypertrophy. The cells were divided into a group with the addition of 5mM beta-hydroxybutyrate (BHB) and a control group (treated with PBS) (Figure 6a). Comparing the cell size after 48 hours, cardiac cell hypertrophy was suppressed in the group treated with BHB (Figures 6b and 6c).

**Figure 6.**
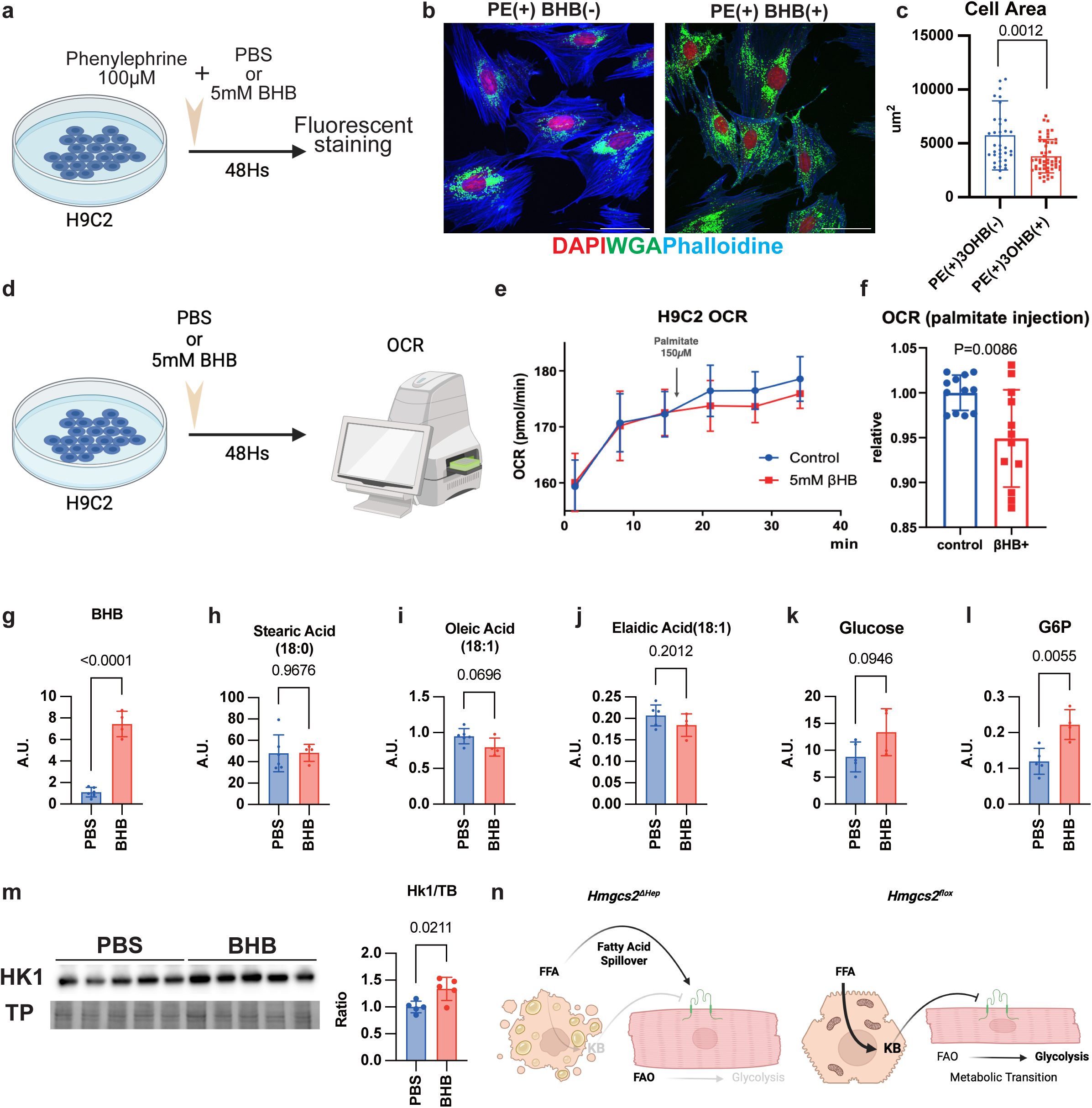
β-Hydroxybutyrate directly suppresses fatty acid utilization and promotes glucose utilization in cardiomyocytes. a, Schema of protocol of inducing hypertrophy for H9C2 cells and measured the cell size. H9C2 cells were induced hypertrophy by 100μM phenylephrine and incubated with or without 5mM β-Hydroxybutyrate (BHB) for 48 hours. b,H9C2 cells immunohistochemical staining with 4’,6-diamidino-2-phenylindole (DAPI), wheat germ agglutinin lectin (WGA), and Phalloidine. Scale bar: 50 μm. c, Area of H9C2 cells d, Schema of incubation and oxygen consumption ratio (OCR) analysis using Seahorse analyzer. H9C2 cells were incubated with or without 5mM BHB for 48 hours, and OCR was measured. e, OCR changes by administration of 150 μM Palmitate incubated with/without 5mM β-HB. f, OCR difference ratio between control and β-HB incubation groups. g-j: concentration of fatty acids and substrates of the glycolytic pathway. g, β-Hydroxybutyrate (BHB). h, Stearic acid. i, Oleic acid. j, Eladic acid. k, Glucose. l, Glucose-6-phosphate (G6P). m, Western blot quantification of Hexokinase 1 (Hk1) protein expression, normalized to total protein (TP). n, Diagram summarizes the results of present research. In Hmgcs2ΔHep, hepatocytes with deficient ketone body synthesis accumulate fatty acids, leading to dysfunction. This results in an oversupply of fatty acids and a lack of ketone bodies to the myocardium, causing metabolism to be dominated by fatty acid oxidation (FAO) (left). On the other hand, in Hmgcs2flox, where ketone body synthesis is preserved, fatty acids are converted into ketone bodies in hepatocytes and supplied to the myocardium, leading to a shift from FAO to glycolysis. c, f-m, Welch’s t-tests were used for statistical analysis.

Following this, to investigate the utilization of the energy substrate, H9C2 cells were cultured with 5mM β-hydroxybutyrate for 48 hours, and oxygen consumption was measured (Figure 6d). While the oxygen consumption rate (OCR) itself showed no significant difference, the increase in oxygen consumption stimulated by 150 μM palmitate was suppressed by the BHB pretreatment (Figures 6e and 6f). This indicated that the utilization of fatty acids as an energy substrate was directly inhibited by prior exposure to BHB. Subsequently, metabolomic analysis of H9C2 cells subjected to similar treatments revealed an expected increase in intracellular BHB levels (Figure 6g) accompanied by a trend of decreased intracellular fatty acid concentrations (Figure 6h-j).

We next turned our attention to the glycolytic pathway. Analysis of key intermediates revealed a trend toward increased intracellular glucose concentration and a significant increase in Glucose-6-Phosphate (G6P) in the BHB pre-treatment group (Figure 6k,l). Furthermore, HK1, enzyme responsible for converting Glucose to G6P, showed increased protein expression in the treatment group (Figure 6m).

Collectively, these *in vitro* results suggest that BHB directly induces a beneficial metabolic shift in cardiomyocytes, characterized by the simultaneously suppressing fatty acid metabolism and promoting glucose utilization.

## Discussion

Our study provides compelling evidence that insufficient hepatic ketogenesis exacerbates cardiac hypertrophy induced by chronic metabolic and hemodynamic stress. We have identified a novel mechanistic cascade where a deficit in liver-derived ketone bodies impairs the heart’s ability to undergo the protective metabolic transition from fatty acid oxidation (FAO) to glycolysis. This failure is characterized by the downregulation of glycolytic enzymes and the sustained dominance of fatty acid metabolism. Our *in vitro* data further confirm that BHB directly acts on cardiomyocytes to suppress FAO and enhance glucose utilization, establishing ketone bodies as critical mediators that coordinate systemic lipid supply with myocardial metabolic responses.

A key finding is the distinct roles of systemic versus local ketogenesis. While myocardial *Hmgcs2* is essential for acute ischemia-reperfusion recovery^34^, its role may be context-dependent. Specially, intrinsic ketogenesis is vital for neonatal mitochondrial maturation and metabolic reprogramming^40^, suggesting the heart’s reliance on self-produced ketogenesis is highest during development or acute crises. In our chronic HFpEF model, however, inducible cardiomyocyte-specific *Hmgcs2* deletion yielded no overt function or structural changes. This is remarkably consistent with the recent findings by Koay et al., who also utilized adult inducible *Hmgcs2* knockout mice and found preserved cardiac structure despite subtle strain alterations in HFpEF^41^. These consistent observations suggest a functional hierarchy: intrinsic cardiac ketogenesis contributes during development, acute stress, or specific pharmacological response like NAD+ therapy, but is less consequential for preventing structural remodeling during chronic HFpEF progression. Instead, the systemic endocrine supply of ketone bodies from the liver serves as the dominant regulator of myocardial metabolic resilience and structural protection.

We propose that BHB serves as a requisite signal for the "fetal gene reprogramming" of cardiac metabolism.^42^ During hypertrophy, the shift toward glucose utilization is typically protective, yet this process is hindered in our *Hmgcs2^ΔHep^* mice. Specifically, we observed an upregulation of PDK4—a potent inhibitor of the pyruvate dehydrogenase complex—and a downregulation of hexokinase (HK). PDK4 levels correlate with the severity of human hypertrophic cardiomyopathy, and its inhibition is known to be cardioprotective.^43,44^ Our findings link systemic ketone deficiency to this protective axis: BHB signaling appears necessary to dampen PDK4 expression and facilitate HK-mediated glucose utilization. Without this "ketone-driven switch," the heart remains locked in a state of FAO dominance, which is maladaptive under chronic stress.

Our results offer a molecular foundation for the emerging Cardiovascular-Kidney-Metabolic (CKM) syndrome,^1^ strongly advocating for the inclusion of the liver ("Hepatic") as a core component—effectively a "CHKM" pathology. While MASLD is known to impair hepatic ketogenesis via *Hmgcs2* downregulation, ^46^ its direct role in driving cardiac dysfunction has been overlooked. We characterize this as a "metabolic mismatch": the failing heart demands alternative fuels and protective signals that the compromised liver fails to provide. This identifies impaired hepatic ketogenesis as a critical, underappreciated link in the Hepato-Cardio Axis.

These insights open new therapeutic avenues. The clinical benefit of exogenous ketone esters in HFpEF patients ^33^ may be explained by their ability to bypass a dysfunctional liver and directly activate myocardial metabolic reprogramming (PDK4 suppression and HK activation). Consequently, therapeutic strategies that improve hepatic mitochondrial function and ketogenesis may exert pleiotropic cardioprotective effects, emphasizing the need for an integrated multisystem approach to CKM syndrome.

### Limitation

We acknowledge several limitations. First, while the metabolic pathways are conserved, the translational validity of this liver–heart axis requires confirmation in human cohorts with metabolic dysfunction. Second, our findings are based on a specific model of combined stress; the role of this metabolic failure in other forms of heart failure, such as pure hypertension or myocardial infarction, remains to be determined. Third, our *in vitro* mechanisms were primarily explored in H9C2 cells; future studies using primary adult cardiomyocytes would further validate BHB’s effects on mature oxidative metabolism. Finally, the precise molecular sensor or signaling cascade by which BHB modulates PDK4 and HK expression remains to be elucidated.

## Conclusion

In conclusion, our study demonstrates that hepatic ketogenesis is not merely a fuel supply system but a critical regulator of cardiac metabolic plasticity. The loss of this liver-heart communication exacerbates pathological remodeling under chronic stress. Targeting this metabolic axis offers a promising therapeutic strategy for patients with heart failure and concurrent metabolic liver disease.

## Acknowledgement

The authors thank Megumi Nagahiro, Saeko Tokunaga, Chiharu Esumi and Yasuyo Kimura for their excellent technical support throughout the experiments. The Kumamoto University School of Medicine Core Laboratory for Medical Research and Education provided support for GC-MS, and microscopic analysis.

